# Effect of Nonlinear Hyperelastic Property of Arterial Tissues on the Pulse Wave Velocity based on the Unified-Fiber-Distribution (UFD) Model

**DOI:** 10.1101/2022.09.27.509711

**Authors:** Hai Dong, Minliang Liu, Julia Woodall, Bradley Leshnower, Rudolph L. Gleason

## Abstract

Pulse wave velocity (PWV) is a key, independent risk factor for future cardiovascular events. The Moens-Korteweg equation describes the relation between PWV and the stiffness of arterial tissue with an assumption of isotopic linear elastic property of the arterial wall. However, the arterial tissue exhibits highly nonlinear and anisotropic mechanical behaviors. There is a limited study regarding the effect of arterial nonlinear and anisotropic properties on the PWV. In this study, we investigated the impact of the arterial nonlinear hyperelastic properties on the PWV, based on our recently developed unified-fiber-distribution (UFD) model. The UFD model considers the fibers (embedded in the matrix of the tissue) as a unified distribution, which expects to be more physically consistent with the real fiber distribution than existing models that separate the fiber distribution into two/several fiber families. With the UFD model, we fitted the measured relation between the PWV and blood pressure which obtained a good accuracy. We also modeled the aging effect on the PWV based on observations that the stiffening of arterial tissue increases with aging, and the results agree well with experimental data. In addition, we did parameter studies on the dependence of the PWV on the arterial properties of fiber initial stiffness, fiber distribution, and matrix stiffness. The results indicate the PWV increases with increasing overall fiber component in the circumferential direction. The dependences of the PWV on the fiber initial stiffness, and matrix stiffness are not monotonic and change with different blood pressure. The results of this study could provide new insights into arterial property changes and disease information from the clinical measured PWV data.

## 1. Introduction

Pulse wave velocity (PWV) is the speed in which the blood pressure pulse propagates through the vascular tree [1]. PWV is a non-invasive, clinically-tractable metric used to assess arterial stiffness. The measurement of carotid to femoral PWV (cfPWV) has been suggested as a tool for assessment of subclinical target organ damage by European Society of Hypertension/European Society of Cardiology guidelines for the management of arterial hypertension [2]. Moreover, PWV has many applications in prediction of cardiovascular diseases. For instance, Vlachopoulos et al. [3] showed that aortic PWV is a strong predictor of future cardiovascular events and all-cause mortality. The study by Yamashina et al. [4] indicates that the brachial-ankle PWV can be served as an indicator of atherosclerotic cardiovascular risk and severity of atherosclerotic vascular damage.

PWV in the artery is highly related with the material properties of the arterial tissue. The well-known Moens-Korteweg equation [5] characterizes the relation between PWV and aortic stiffness, i.e., 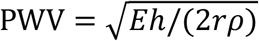, where *E* is the Young’s modulus, *h* is the wall thickness, and *r* is the radius of the artery, and *ρ* is the blood density. The Moens-Korteweg equation assumes an isotopic linear elastic property of the arterial tissue. However, the arterial tissue exhibits highly nonlinear and anisotropic mechanical behaviors [6-10]. Thus, the Moens-Korteweg equation does not capture the relation between PWV and the properties of nonlinearity and anisotropy.

The arterial tissue is comprised of networks of collagen fibers embedded in a ground matrix, which can be regarded as fiber reinforced composites. The tissue is usually stiffer in the circumferential direction than in the axial direction [11], and the degree of anisotropy of arteries is determined primarily by the collagen fiber distribution. The nonlinear mechanical response of the arterial tissue is attributed to progressive engagement of collagen fibers under increases in strain. The fiber distribution and stiffening effect will affect the PWV in the artery. Few existing studies have investigated the impact of material non-linearity and anisotropy on the PWV [12, 13].

In this study, we analyzed the effect of hyperelastic properties of arterial tissues on the PWV. Specifically, we employed the unified-fiber-distribution (UFD) model to model the non-linear, anisotropic, hyperelastic response of the aorta and characterized the dependence of the PWV on the fiber distribution, fiber stiffening effect, fiber initial stiffness and the matrix stiffness. We also modeled the evolution of PWV with aging based on observations that aging induces arterial stiffening [14, 15]. The results indicate the PWV increases with increasing overall fiber component in the circumferential direction. The dependences of the PWV on the fiber initial stiffness, and matrix stiffness are not monotonically and change with different blood pressure. The results of this study could provide new insights into aortic disease information reflected by clinical measured PWV data. The relation between the PWV and the hyperelastic properties may also be applied to obtain the hyperelastic parameters by inverse methods.

## 2. Methods

### 2.1 Unified-Fiber-Distribution (UFD) Hyperealstic Model

Large arteries are composed of three layers, the intima, media and adventitia layers [16]. The tissue of each layer is comprised of networks of collagen fibers embedded in a ground matrix and can be modeled as fiber reinforced composites, of which the mechanical properties are anisotropic [9, 17, 18], similar to engineering fiber-reinforced composites [19-26]. Schriefl at al. [27] showed that there are significant dispersions for the collagen fiber distributions in the three layers of human arteries, and each layer may contain multiple fiber families. Their study [27] also revealed that the number of fiber families may vary from different positions for the same kind of tissue, such as there are two fiber families in the media of human descending thoracic aorta and abdominal aorta, while there is only one fiber family in the media of common iliac artery. In the widely used Gasser-Ogden-Holzapfel (GOH) model [28], a fiber distribution was separated into two fiber families. Each of the fiber families is described by a transversely isotropic von Mises distribution. The assumption of two fiber families with transversely isotropic von Mises distribution may be not physically consistent with the real fiber distribution of aortic tissues. The unified-fiber-distribution (UFD) model [29, 30] considers the fibers as a unified distribution, rather than separating the fiber distribution into two/several fiber families. The consideration of a unified distribution for the fibers in the UFD model [29-32] may be more physically consistent with the real fiber distribution of aortic tissues.

Arterial tissues are considered to be incompressible, non-linear, elastic, homogenous, and anisotropic and experience large strains under physiologically-relevant loads. The strain energy function of the UFD model can be expressed as

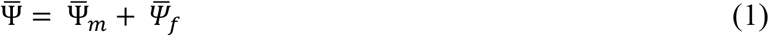

where 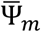 is the strain energy of the matrix contribution given by the neo-Hookean model [33-37]

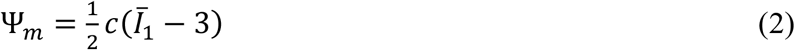

where *c* is the initial shear modulus of the matrix, 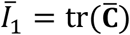 is the first invariant of 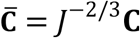, with **C** = **F**^T^**F** denoting the right Cauchy–Green tensor, in general, *J* = det **F** > 0, where **F** is the deformation gradient. We have *J* ≡ 1 since the tissues are considered to be incompressible. The 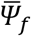 is the strain energy of the fiber contribution which distinguishes the UFD model from other models, given by [29, 30]

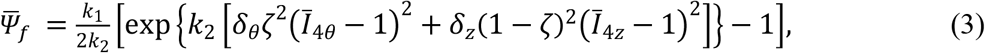

where 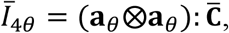, and 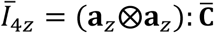 with **a**_***θ***_ and **a**_***z***_ denoting the unit vectors in the circumferential direction and the axial direction in the reference basis, respectively. The roles of *δ*_***θ***_ and *δ*_*z*_ are to exclude the contribution of the compressed fibers, given by the Heaviside step function

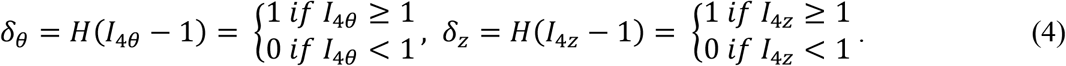

The parameters *k*_1_, *k*_2_ and ζ in Eq. (3) are fiber related material parameters. The parameter ζ ∈ [0,1J represents the average fiber component in the circumferential (**a**_***θ***_) direction, and (1 - ζ) represents the average fiber component in the axial (**a**_*z*_) direction. ζ = 1 corresponds to the distribution of all the fibers along the circumferential direction, and ζ = 0 represents the distribution of all the fibers along the axial direction. To illustrate how the parameter ζ affects the mechanical behavior of arterial tissue, we consider an in-plane equal biaxial stretch (thickness direction stress-free), with *λ*_*θ*_ = *λ*_*z*_ = *λ* > 1, where *λ*_*θ*_ and *λ*_*z*_ are stretches in the circumferential and axial directions, respectively. The component of the Cauchy stress *σ*_*θθ*_ and σ_*zz*_ can be expressed as (see Eq. (A5) in Appendix A, with *λ*_*θ*_ = *λ*_*z*_ = *λ* and *δ*_*θ*_ = *δ*_*z*_ = 1)

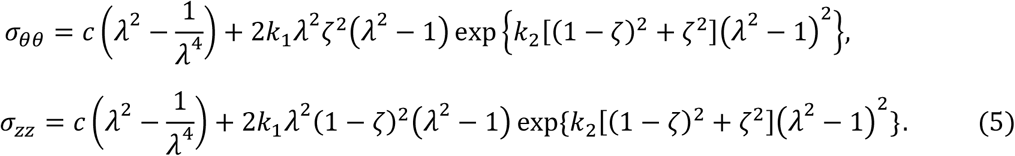

where the term 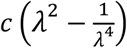 is the contribution of the matrix, and the term with *k, k* and ζ is the contribution of the fiber, noted by

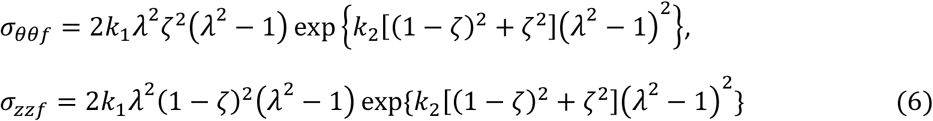

With fixed values for *k*_1_ and *k*_2_, based on Eq. (5), the tissue/fiber stiffness increases in the circumferential direction (Fig. 1a), and decreases in the axial direction with increasing ζ (Fig. 1b).

**Fig 1.**
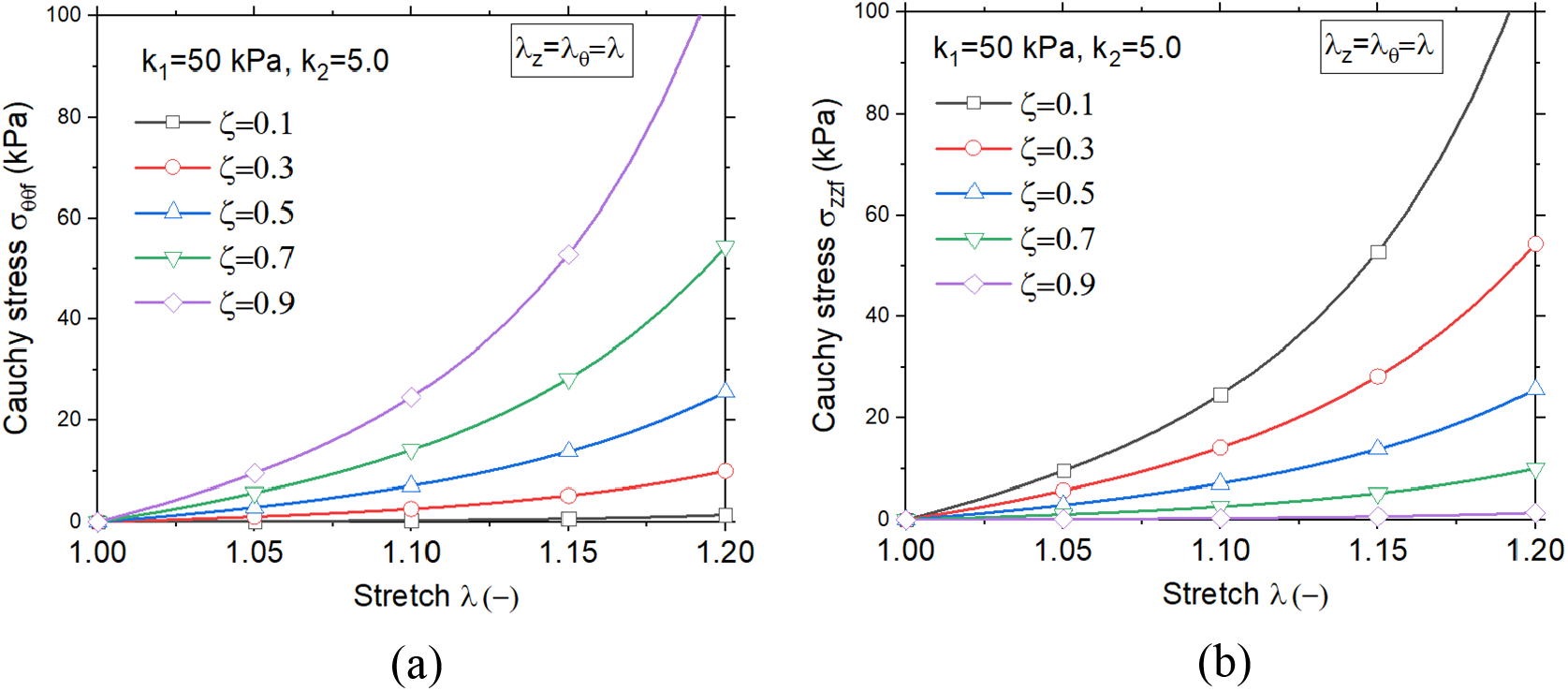
Fiber contribution stress-stretch relation under in-plane equal biaxial stretch (*λ*_*θ*_ = *λ*_*z*_ = *λ*): (a) *σ*_*θθf*_ vs. *λ*: stiffness increases in the circumferential direction with increasing ζ; (b) *σ*_*zzf*_ vs. *λ*: stiffness decreases in the axial direction with increasing ζ.

To illustrate the physical meanings of *k*_1_ and *k*_2_, we plotted (Fig. 2) the dependence of σ_00f_ on *k*_1_ and *k*_2_ based on Eq. (5)_1_, with fixed value of ζ = 0.5. The initial slope (i.e., initial modulus) of the fiber stress-stretch (*σ*_*θθf*_ - *λ*_*θ*_) curve increases proportionally with increasing *k*_1_, while the stiffening effect, characterized by the ratio between the stiffening tangent modulus and the initial tangent modulus, does not change (Fig. 2). For the fiber stress-stretch (*σ*_*θθf*_ - *λ*_*θ*_) curve for different *k*_2_ with fixed *k*_1_ the initial slope (initial modulus) does not change, while the stiffening effect increases with increasing *k*_2_. Similar results could be obtained for the fiber stress-stretch (*σ*_*θθf*_ - *λ*_*θ*_) curve. Thus, the *k*_1_ parameter controls the initial modulus of the fiber and *k*_2_ controls the stiffening effect.

**Fig 2.**
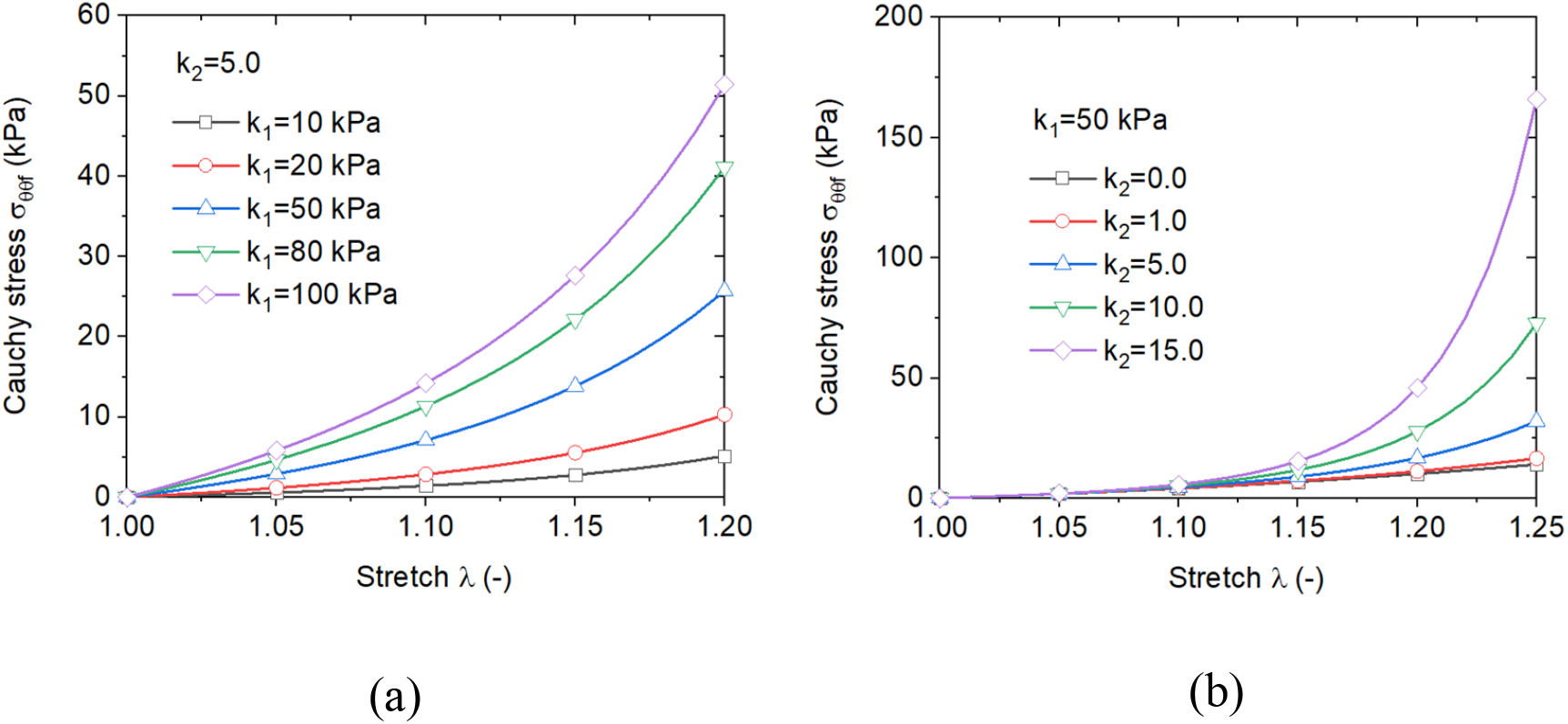
Fiber stress-stretch relation: (a) changed *k*_1_ with fixed *k*_2_, the initial slope (initial modulus) of the curve increases proportionally with increasing *k*_1_, while the stiffening effect does not change; (b) changed *k*_2_ with fixed *k*_1_, the initial slope of the curve does not change, while the stiffening effect increases with increasing *k*_2_.

### 2.2 Pulse Wave Velocity (PWV) in a Hyperelastic Artery Tube

#### 2.2.1 Relation between PWV and blood pressure

Here, we consider the case of nonviscous blood flowing in a cylindrical hyperelastic arterial tube without wave reflection. The PWV can be expressed as [5]

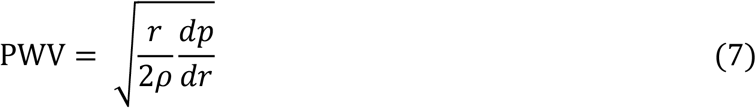

where *p* is the inside blood pressure, *ρ* is the density of the blood flow, and *r* is the radius of the artery tube in the deformed configuration under blood pressure *p*.

#### 2.2.2 Relation between PWV and the 2nd P-K stress

Based on the Laplace law, we have

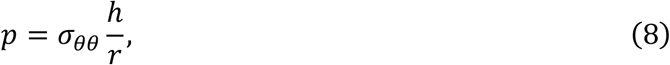

where *σ*_*θθ*_ is the Cauchy stress in the circumferential direction of the artery, and *h* is the thickness of the artery tube in deformed configuration. The parameters *r* and *h* can be further expressed as

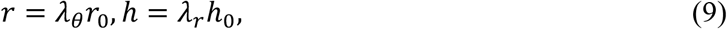

where *r*_0_ and *h*_0_ are the initial radius and thickness, respectively, in the reference/undeformed configuration. The parameters *λ*_*θ*_ and *λ*_r_ are the stretch in the circumferential and radius directions, respectively.

The deformation gradient of the cylindrical artery tube under inner pressure can be expressed as **F** = diag[*λ*_r_, *λ*_*θ*_, *λ*_*z*_] = *diag*[(*h*/*h*_*o*_), (*r*/*r*_*o*_), (*λ*/*λ*_*o*_)J, where *h, r*, and *λ* and *h*_*o*_, *r*_*o*_, and *λ*_*o*_ are the vessel thickness, radius, and axial length in the loaded (in vivo) and unloaded configurations, respectively (Fig. 3). With the assumption of incompressibility, we have det**F** = *λ*_*r*_*λ*_*θ*_*λ*_*z*_ = 1, and further we have

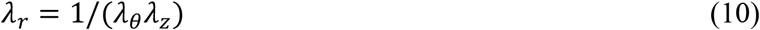

Using the relation between Cauchy stress **σ** and the 2nd PK stress **S** (i.e. σ = **FSF**^*T*^) we have

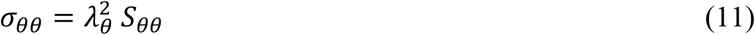

With the combination of Eqs. (8)-(11), we have

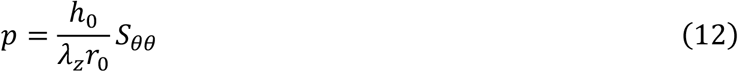

Here, we assume the pre-stretch *λ*_*z*_ is fixed during the pressure wave propagation and we further have

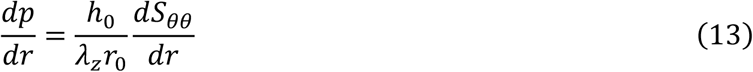

With Eq. (9)_1_, we have

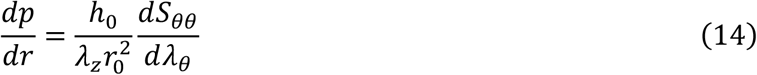

Substituting Eq. (14) into (7) yields

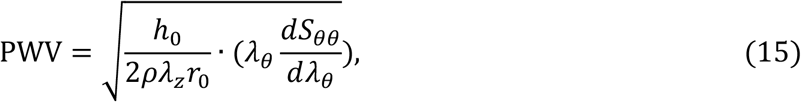

**Fig 3.**
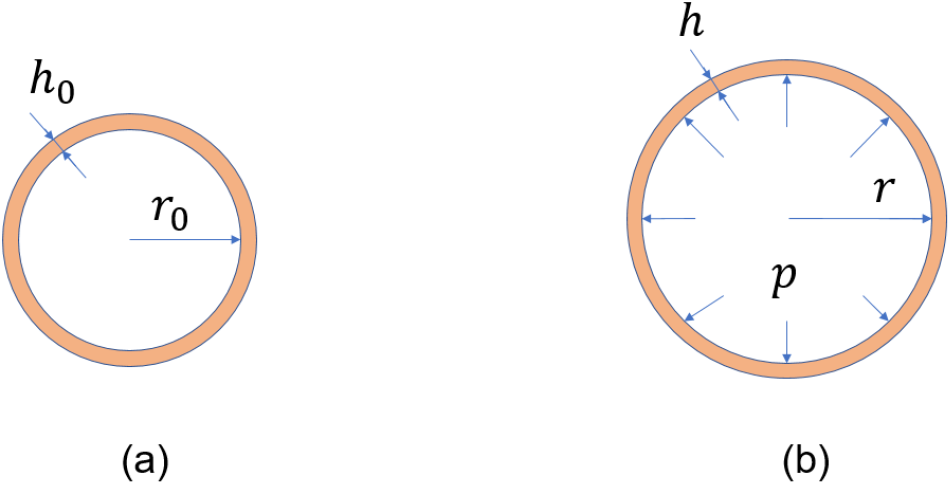
Schematic of the cross-section of a cylindrical artery tube: (a) reference configuration with un-deformed radius *r*_0_ and thickness *h*_0_; (b) current configuration under inner pressure *p* with deformed radius *r* and thickness *h*.

#### 2.2.3 Relation between PWV and hyperelastic parameters

For the cylindrical artery tube under inner pressure with fixed pre-stretch in the axial direction, *S*_*θθ*_ can be expressed as (see Eq. (A6) in Appendix A),

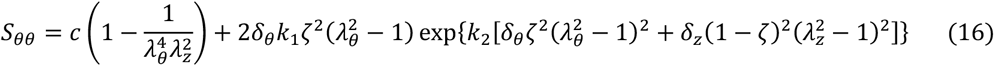

Further we have,

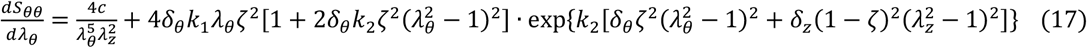

Substituting Eq. (17) into (15) yields

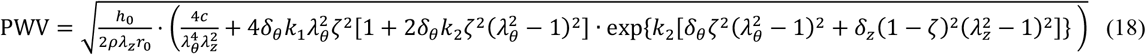

Equation (18) can be represented as a function as

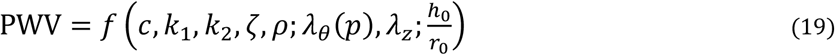

where *λ*_*θ*_ is uniquely determined by the blood pressure *p* when other parameters is fixed. Equations (18) and (19) indicate that the value of PWV is affected by the arterial and blood flow properties { *c, k*_1_, *k*_2_, ζ, *ρ*}, the loading conditions {*p, λ*_*z*_}, and the un-deformed geometric parameter ratio 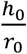.

## 3. Results

### 3.1 Fitting of Experimentally Measured Data

#### 3.1.1 Relation between blood pressure (BP) and PWV

As shown in Eqs. (18)/(19), the blood pressure (*p*) influences the value of PWV. Chen et al. [38] experimentally observed this by measuirng the ear-to-toe PWV at different blood pressures in a single subject. We fitted the experimental data of pressure-PWV relation based on Eq. (18), and good fitting results were obtained (Fig. 4), with *R*^2^ = 0.9307 and root-mean-square-error, RMSE=1.23 m/s. During the fitting, we assumed *h*_*o*_/*r*_*o*_ = 0.15 [13], *ρ* = 1.056 g · cm^-3^, and *λ*_*z*_ = 1.05. The fitted material parameters are *c* = 405.05 kPa, *k*_1_ = 9.19 kPa, *k*_2_ = 325.55, and ζ = 0.9823. We noted that there are some discrepancies within the lower range of blood pressure (∼50-70 mmHg). The reason could be that we assumed a fixed value for the axial pre-stretch (*λ*_*z*_) which may vary with changing blood pressure. This was investigated with a parameter study in Sec. 3.2.

**Fig 4.**
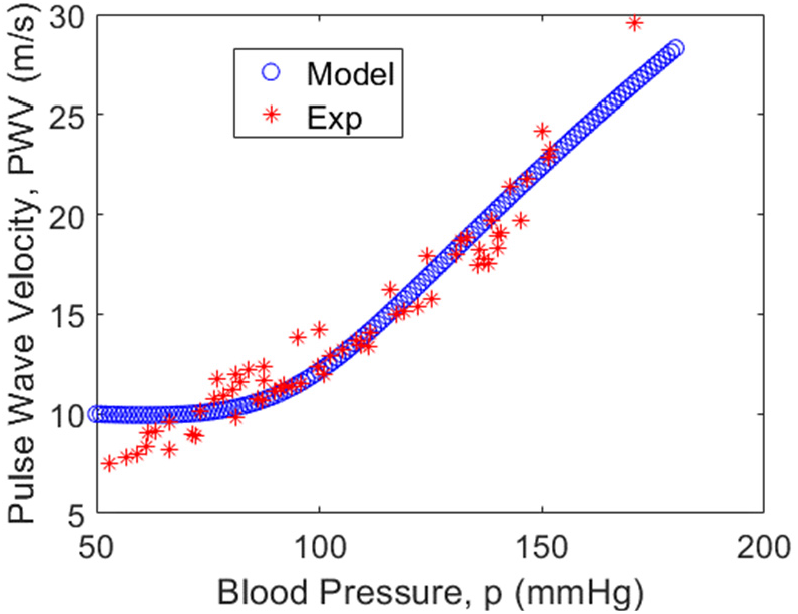
Comparison between theoretical (Model) and experimental (Exp) results for the p-PWV relation.

#### 3.1.2 Effect of age-induced stiffening on PWV

The stiffness of human arterial tissue could increase with aging [15], which further results in the increase of the PWV. Rogers et al. [39] measured the aortic PWV of healthy human with different ages. They found that the PWV from thoracic ascending to thoracic descending aorta significantly increases with growing age (red stars in Fig. 5). Ferruzzi et al. [14] showed that the wavelength of curved fibers (Fig. 3 in Ref [14]) in the aortic tissue of mice significantly increases with aging (from 20 to 100 weeks old), which could affect the stiffening effect of the tissue under larger stress loading. Recall that the parameter *k*_2_ in Eq. (3) represents the stiffening effect. To characterize the age-induced stiffening, here we assume a simple linear relation between *k*_2_ and the age, i.e.,

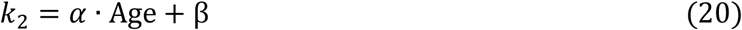

With Eq. (20), Eq. (19) can be re-formed as

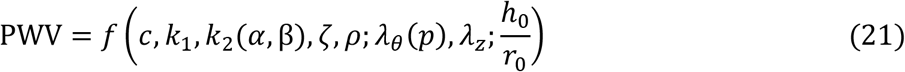

We fitted the experimental data from Rogers et al. [39] with PWV-Age (Fig. 5) based on Eq. (21) and obtained good fitting results, with *R*^2^ = 0.6532 and RMSE=1.76 m/s. During the fitting, we assumed *h*_0_/*r*_0_ = 0.15 [13], *ρ* = 1.056 g · cm^-3^, and *λ*_*z*_ = 1.2. For simplicity, we also fixed the value of β = 0 to reduce the degree of over-parameterization. The fitted parameters are *c* = 124.78 kPa, *k*_1_ = 7.39 Pa, *α* = 0.0253/year, and ζ = 0.3893. During the fitting, we have assumed only the parameter *k*_2_ changes with age, and the parameters *c, k*_1_, and ζ keep the same. However, these values may also change during growth with age. Moreover, the values of axial pre-stretch (*λ*_*z*_) and thickness-radius ratio could also be different for different subjects. Nevertheless, the *in vivo* measurement of their influence on PWV is difficult. We will do parameter studies on how the change of these parameters affects the PWV in the following section.

**Fig. 5.**
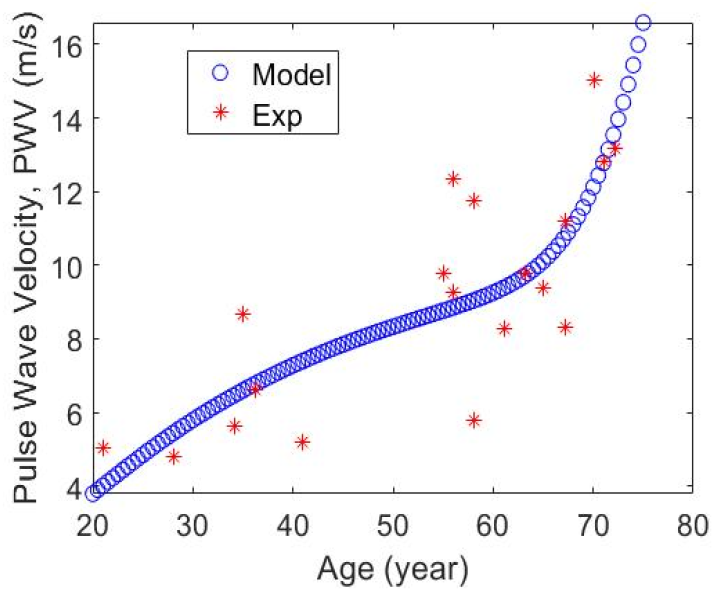
Comparison between theoretical (Model) and experimental (Exp) results for the Age-PWV relation.

### 3.2 Parameters Studies

#### 3.2.1 Impact of fiber distribution (ζ), fiber initial stiffness (*k*_1_), and matrix stiffness (*c*)

The PWV increases with increasing fiber initial stiffness (*k*_1_) for relatively low pressure (e.g., 50 mmHg), while decreases with increasing fiber initial stiffness (*k*_1_) for relatively high pressure (Fig. 6a). The value of circumferential stretch *λ*_*θ*_ reduces with increasing *k*_1_ when the blood pressure is fixed (see Fig. 6b), which tends to decrease the value of PWV (see Eq. (15)). For low blood pressure (e.g., 50 mmHg), the value of *λ*_*θ*_ is relatively small (Fig. 6b) and the trend of PWV in Fig. 6a is dominated by the increasing stiffness (*k*_1_) which results in the increase of PWV with increasing *k*_1_. For relatively high blood pressure (e.g., 160 mmHg), the value of *λ*_*θ*_ is large (Fig. 6b) and the trend of PWV in Fig. 6a is dominated by the changing *λ*_*θ*_, which results in the decrease of PWV with increasing *k*_1_. The baseline values of the parameters in Fig. 6 and the following Figs. 7-8 are *c* = 100 kPa, *k*_1_ = 20 kPa (not for Fig. 6), *k*_2_ = 100, ζ = 0.7, *ρ* = 1.056 g · cm^-3^, *λ*_*z*_ = 1.05, *h*_0_/*r*_0_ = 0.15.

**Fig 6.**
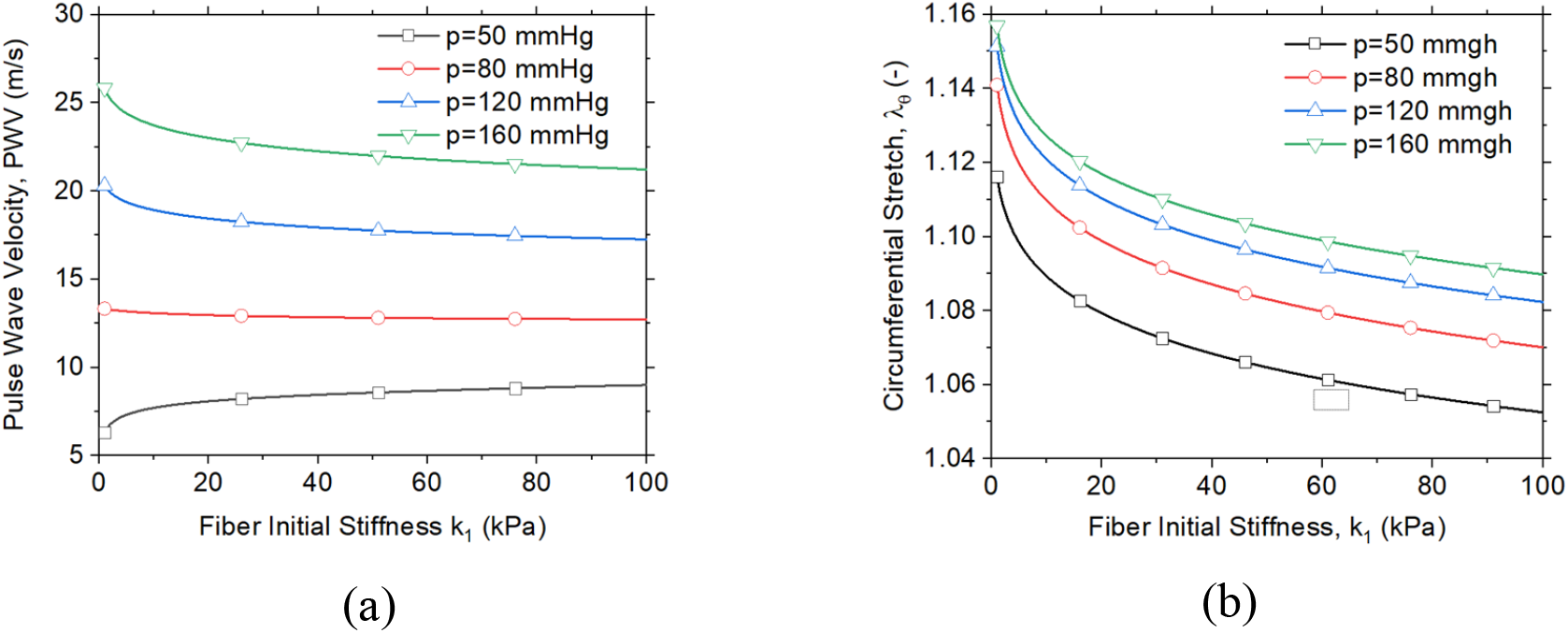
(a) Relation between the PWV and fiber initial stiffness (*k*_1_) with different blood pressure (*p*); (b) Relation between circumferential stretch *λ*_*θ*_ and the fiber initial stiffness (*k*_1_), with different blood pressure.

**Fig 7.**
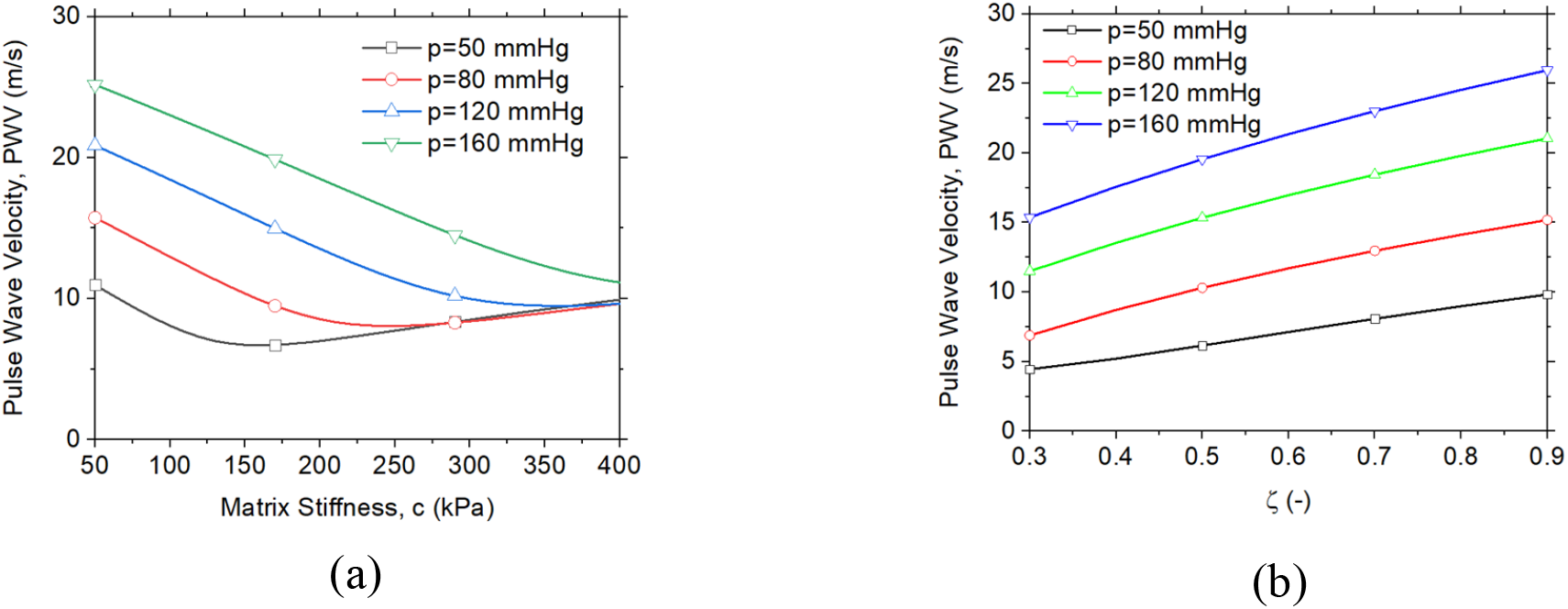
(a) Relation between the PWV and the matrix stiffness under different blood pressure (*p*); (b) Relation between the PWV and circumferential fiber component (ζ) under different blood pressure (*p*).

**Fig 8.**
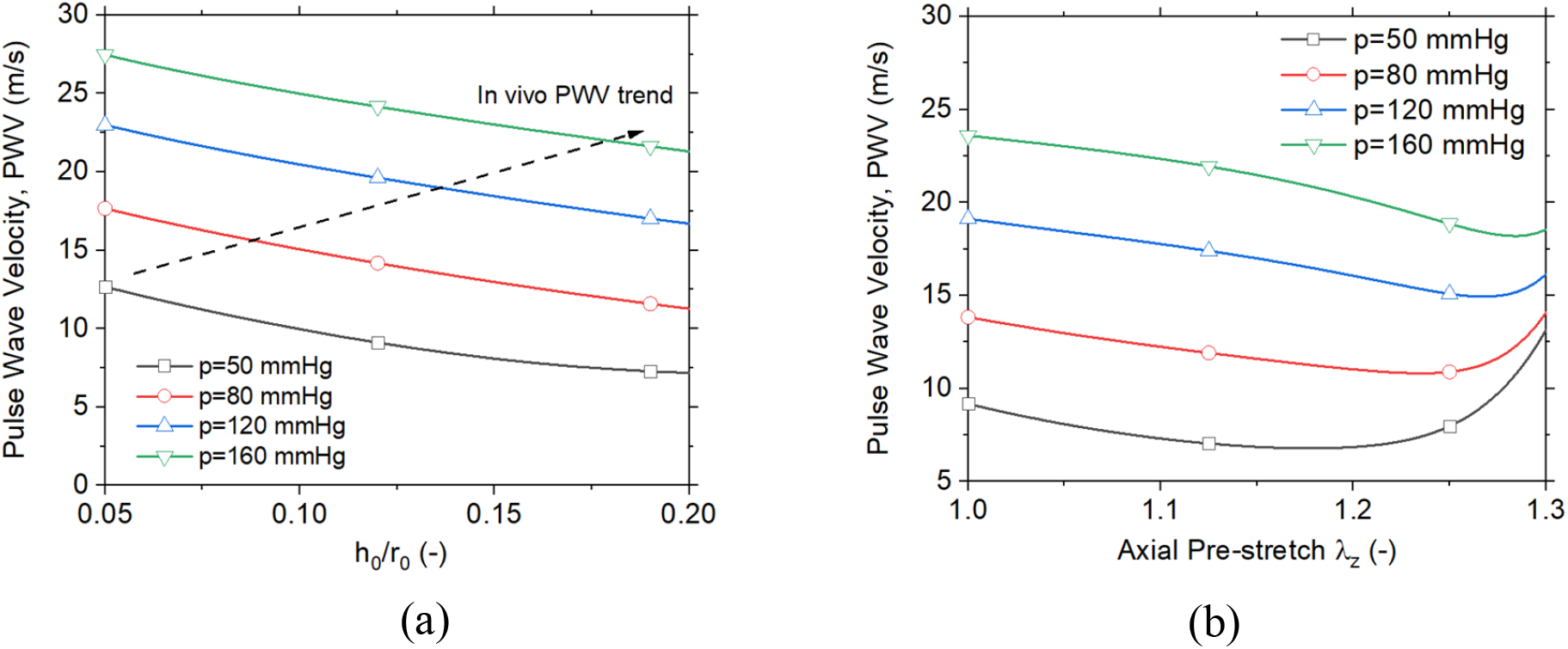
(a) Relation between the PWV and the undeformed radius *r*_0_. (b) Relation between the PWV and the undeformed thickness *h*_0_.

As to the impact of matrix stiffness in Fig. 7a, for low blood pressure (e.g., 50 mmHg), PWV first decreases and then increases with increasing matrix stiffness. For high blood pressure (e.g., 160 mmHg), PWV decreases monotonically with increasing matrix stiffness. Recall that ζ represents the average fiber component in the circumferential, and the circumferential stiffness increases with increasing ζ, which results in the rise of the PWV with increasing ζ (Fig 7b).

#### 3.2.2 Impact of thickness-radius ratio and axial pre-stretch

The PWV decreases with increasing thickness-radius ratio when the blood pressure is fixed (Fig. 8a). Increasing thickness-radius ratio results in the rise of structure stiffness of artery. However, with the same blood pressure, the circumferential stretch *λ*_*θ*_ also decreases with the thickness-radius ratio, which results in the decline of PWV. For *in vivo* cases, thickening of artery walls usually also raises the blood pressure, and thus the black dash line could be the real dependent path of PWV on the thickness-radius ratio. The PWV first decreases and then increases with increasing axial pre-stretch (Fig. 8b). This complex relation between PWV and axial pre-stretch also resulted from the coupled influence of *λ*_*z*_ on the circumferential stretch *λ*_*θ*_.

## 4. Discussions and Conclusions

The dependence of PWV on mechanical properties of arterial tissue is complex, due to the coupling influences of stiffness and deformation on PWV. The simple Moens-Korteweg equation [5] is not enough to characterize relation between PWV and the arterial properties with nonlinearity and anisotropy. For instance, the PWV could possible decreases with increasing arterial/fiber initial stiffness for certain blood pressures (Fig. 6a). Similar results were observed by Milkovich et al. [40] which shows that the PWV could possibly be negatively related to the increasing modulus, based on experimental measurements. Such kind of negative dependence of PWV on modulus can be hardly explained based on the Moens-Korteweg (MK) equation, even with the improvement of Hughes equation [13, 41] which assumes the Young’s modulus in MK equation increases with blood pressure.

In general, there are significant subject-specific variations in the mechanical properties of the arterial/aortic wall. It is still a challenging problem to accurately identify the *in vivo* nonlinear and anisotropic properties of arterial/aortic walls. Based on clinical images (e.g., CT data), traditional inverse methods via finite element (FE) simulation could calculate the *in vivo* mechanical properties [42-45]. However such methods are usually computationally expensive, taking from several hours to weeks [42]. Here in Section 3.1.1, we calculated the *in vivo* mechanical properties of arterial wall with optimization based on the PWV data at different blood pressures (Fig. 4), which only takes a few minutes. Such kind of PWV-based inverse method may be a promising way to fast identify the *in vivo* nonlinear and anisotropic properties of arterial/aortic wall.

The accuracy in fitting of PWV-age relation (Fig. 5) is not as good as that of PWV-p relation (Fig. 4). The possible reason could be that the PWV-p data in Fig. 4 was obtained based on measurements of a single person [38] while the PWV-age in (Fig. 5) were measured with multiple subjects at different ages. The variations of mechanical properties of aortic wall in different individuals could result in the scatter of the PWV-age data points along the fitting curve.

There are several limitations of this study. Firstly, we considered the artery as a perfect cylindrical tube which may be different from the geometry of real patient arteries. Patient-specific geometry may be obtained from clinical CT images [46]. Secondly, The stiffening of human arterial tissue with aging could result from the increased deposition, cross-linking, and micro-structural change of collagen/elastin fibers [14, 15]. Thus, the stiffening with aging could be complicated and nonlinear. In this study, we applied a simple linear relation (Eq. (20)) to characterize the arterial stiffening during aging, which may not reflect the actual stiffening evolution process. Finally, the blood flow was assumed as one-dimensional homogeneous flow, and the reflection was not considered. The *in vivo* flow may be inhomogeneous and the fluid-structure interaction (FSI) simulations [47-53] may be applied to model the three-dimensional flow field.

In conclusion, we investigated the effect of the arterial nonlinear hyperelastic properties on the PWV, based on our recently developed unified-fiber-distribution (UFD) model for arterial tissues. Specifically, we fitted the measured relation between the PWV and blood pressure which obtained a good accuracy. We also modeled the aging effect on the PWV based observations [14, 15] that the stiffening of arterial tissue increases with aging, and the results agrees well with experimental data. In addition, we did parameter studies on the dependence of the PWV on the arterial properties of fiber distribution, fiber initial stiffness, and matrix stiffness. The results indicate the PWV increases with increasing overall fiber component in the circumferential direction. The dependences of the PWV on the fiber initial stiffness, and matrix stiffness are not monotonically and change with different blood pressure, which resulted from the coupling influences of stiffness and deformation on PWV. The results in this study could provide new insights into arterial property changes and disease information from the clinical measured PWV data.

## Acknowledgment

This study is supported by NIH (R01HL155537 and R01HL142036).

## Appendix A: Stress calculation

The Cauchy stress of the arterial tissue can be expressed as [33]

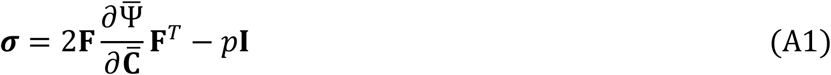

where *p* is a Lagrange contribution to the hydrostatic pressure, and

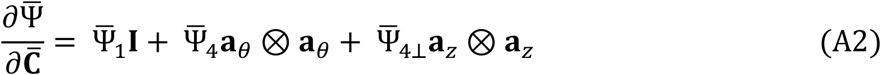

where

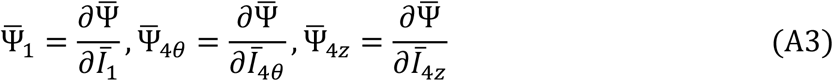

Substituting Eqs. (1)-(3) into Eq. (A3), we have

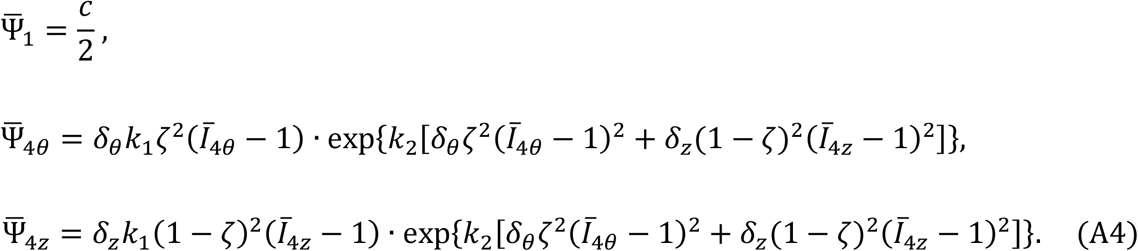

For the cylindrical artery tube under inner pressure with fixed axial pre-stretch, the deformation gradient can be expressed as **F** = diag[1/(*λ*_*θ*_*λ*_*z*_), *λ*_*θ*_, *λ*_*z*_]. Then we have 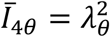 and 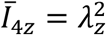. Substituting *Ī*40 and *Ī*4z into Eq. (A4) and then into Eqs. (A1)-(A3), together with the condition of *σ*_*rr*_ = 0, the non-zero components of the Cauchy stress can be expressed as

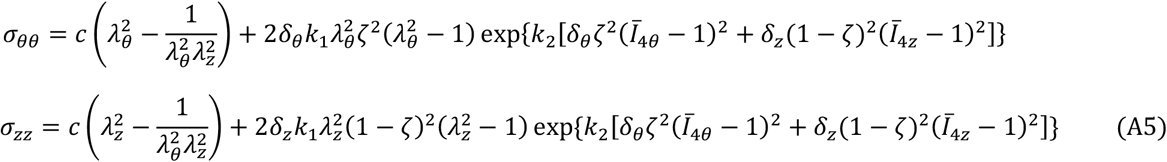

Further, the non-zero components of the 2nd P–K stress can be expressed as

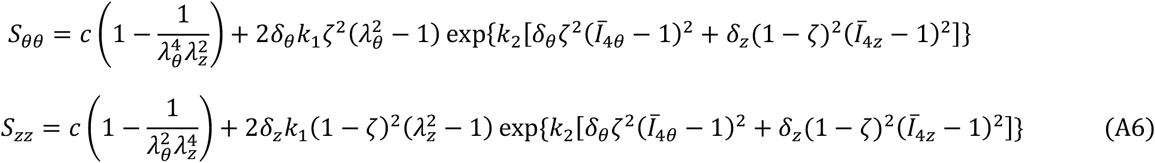

## Notes

### Competing Interest Statement

The authors have declared no competing interest.

